# Automating Candidate Gene Prioritization with Large Language Models: Development and Benchmarking of an API-Driven Workflow Leveraging GPT-4

**DOI:** 10.1101/2024.12.10.627808

**Authors:** Taushif Khan, Mohammed Toufiq, Marina Yurieva, Nitaya Indrawattana, Akanitt Jittmittraphap, Nathamon Kosoltanapiwat, Pornpan Pumirat, Passanesh Sukphopetch, Muthita Vanaporn, Karolina Palucka, Basirudeen Kabeer, Darawan Rinchai, Damien Chaussabel

## Abstract

In this exploratory study, we developed an automated workflow that leverages Large Language Models, specifically GPT-4, to prioritize candidate genes for targeted assay development. The workflow automates interaction with OpenAI models and enables prompt creation, submission. It features customizable prompts designed to evaluate candidate genes based on criteria such as association with biological processes, biomarker potential, and therapeutic implications, which can be tailored for specific diseases or processes. Benchmarking experiments comparing the performance of the Application Programming Interface (API)-based automated prompting approach with manual prompting demonstrated high consistency and reproducibility in gene prioritization results. The automated method exhibited scalability by successfully prioritizing genes relevant to sepsis from the BloodGen3 repertoire, comprising 11,465 genes, distributed among 382 modules. The workflow efficiently identified sepsis-associated genes across the repertoire, revealing distinct gene clusters and providing insights into their distribution within module aggregates and individual modules. This proof-of-concept study demonstrates how LLMs can enhance gene prioritization, streamlining the identification process for targeted assays across various biological contexts. However, it also reveals the need for further validation and highlights the exploratory nature of this work due to scoring inconsistencies and the necessity for manual fact-checking. Despite these challenges, the automated workflow holds promise for accelerating targeted assay development for disease management and paves the way for future research.

## INTRODUCTION

Candidate gene prioritization plays a crucial role in identifying potential biomarkers from large-scale molecular profiling data. Systems-scale profiling technologies, such as transcriptomics, have revolutionized biomedical research by simultaneously measuring tens of thousands of analytes, leading to significant advances in various medical fields, including oncology (1,2), autoimmunity (3,4), and infectious diseases (5,6). However, translating these findings into actionable clinical insights often requires the identification of relevant analyte panels and the design of targeted profiling assays (7,8). Targeted transcriptional profiling assays enable precise, quantitative assessments of the abundance of panels comprising tens to hundreds of transcripts (9,10). These assays offer several advantages, including cost-effectiveness, rapid turnaround times, and the ability to process large sample numbers, making them valuable for research and clinical applications (11–13). However, the critical task of selecting relevant candidate genes for inclusion in targeted assays can be challenging, especially when contending with the extensive volumes of biomedical information generated by systems-scale profiling technologies.

Knowledge-driven methods for candidate gene prioritization must efficiently sift through vast amounts of literature to identify the most promising candidates. This process can be lengthy and may lack depth due to the sheer volume of information available for each gene (14). While resources such as gene ontologies and curated pathways can help, they often provide only superficial information about the genes and may lack context. As a result, there is a pressing need for more efficient and effective methods to assimilate and synthesize the extensive, context-rich information necessary for effective gene curation and analysis.

The introduction of Large Language Models (LLMs) has opened new possibilities for leveraging collective biomedical knowledge in candidate gene prioritization. LLMs, such as GPT-4 (OpenAI), Claude (Anthropic), and PaLM (Google), have demonstrated remarkable capabilities in natural language understanding and generation (15–17) . These models are trained on vast amounts of text data, allowing them to assimilate and synthesize information from diverse sources, including scientific literature. The potential of LLMs in assisting with candidate gene prioritization has been recently explored. In a proof-of-concept study, we demonstrated the utility of LLMs in a manual candidate gene prioritization workflow (18). The study focused on prioritizing genes forming a circulating erythroid cell blood transcriptional signature, which was previously associated with respiratory syncytial virus (RSV) disease severity (5), vaccine response (6), and elevated transcript abundance in patients with metastatic melanoma (5). This manual workflow involved several tasks, such as identifying functional convergences among candidate genes, scoring genes based on specific criteria (e.g., relevance to erythropoiesis, potential as a biomarker), and selecting top candidates for further characterization. We benchmarked four LLMs (GPT-3.5, GPT-4, Bard, and Claude) across multiple evaluation criteria. GPT-4 and Claude demonstrated superior performance through consistent scoring across multiple iterations (correlation coefficients > 0.8), strong alignment with manual literature curation, and well-reasoned, evidence-based justifications for their assessments. This comprehensive evaluation established GPT-4 and Claude as the most reliable tools for our gene prioritization tasks. The study highlighted the potential of LLMs in enhancing the efficiency and depth of knowledge-driven candidate gene prioritization, with minimal human input.

Building upon this work, we sought to further streamline the candidate gene prioritization process by developing an automated LLM-enabled workflow. The transition from manual prompting via the chat interface to automated prompting using Application Programming Interfaces (APIs) offers several potential advantages. First, API-based automation enables the efficient generation and submission of prompts for a large number of candidate genes, reducing the time and effort required for manual input. Second, it allows for the seamless integration of the LLM-based prioritization workflow with other computational tools and pipelines, facilitating the development of end-to-end solutions for targeted assay design. Third, the API-based approach provides a more standardized and reproducible way of interacting with LLMs, minimizing the variability introduced by human operators. This automated workflow aims to enable the prioritization of extensive module repertoires, such as BloodGen3 (10), and facilitate the design of disease-specific panels, such as ones we previously designed as part of our COVID-19 and pregnancy work (9,19). The integration of LLMs into a partially automated candidate gene prioritization pipeline could thus significantly advance biomedical research by accelerating the translation of systems-scale profiling data into targeted assays for clinical and research applications.

## METHODS

### Leveraging APIs for automated prompt submission in gene prioritization

We developed Python scripts to automate communication with GPT-4 through two different APIs: the OpenAI API and the Microsoft Azure API. These scripts handle prompt generation, API communication, response processing, and data storage. The OpenAI API was accessed using the openai library, while the azure-ai-openai library was used for the Microsoft Azure API.

The use of two APIs provides flexibility, allowing users to choose based on factors such as availability, performance, or organizational preferences. This dual-API approach also ensures redundancy and facilitates potential performance comparisons. Our scripts encapsulate the entire workflow, from prompt generation to data organization, streamlining the automated candidate gene prioritization process.

### Automated candidate gene prioritization

Prompts used for gene scoring and prioritization: To prioritize candidate genes using GPT-4, we designed a set of prompts that elicit specific information about each gene, including its official name, function summary, and scores for various criteria relevant to the gene’s potential as a biomarker or therapeutic target. The prompts were carefully crafted to cover essential aspects such as the gene’s association with different types of interferon responses, relevance to circulating leukocytes immune biology, current use as a biomarker in clinical settings, potential value as a blood transcriptional biomarker, known drug target status, and therapeutic relevance for diseases involving the immune system. The scoring system was designed to provide a quantitative measure of the evidence supporting each criterion, with scores ranging from 0 to 10. A score of 0 indicates no evidence found, while a score of 10 represents strong evidence. Intermediate scores reflect varying levels of evidence quality and reliability, with 1-3 indicating very limited evidence, 4-6 indicating some evidence that needs validation or is limited to certain conditions, and 7-8 indicating good evidence. For each gene, the automated prioritization process generates individual scores ranging from 0 to 10 for each of the eight evaluated criteria (a-h). To obtain a single, comprehensive measure of a gene’s overall relevance and importance, we calculate a cumulative score. This cumulative score is the sum of the individual scores across all eight criteria, resulting in a maximum possible score of 80 (8 criteria * 10 points each). This approach allows for a straightforward ranking of genes based on their overall performance across multiple relevant factors.

### API-based automation of the prompting process

To automate the candidate gene prioritization process, we integrated the designed prompts into the Python scripts that communicate with GPT-4 via the OpenAI and Microsoft Azure APIs. The scripts automatically generate the prompts for each candidate gene by incorporating the gene symbol and the predefined scoring criteria. The generated prompts are then sent to GPT-4 through the respective API, and the model’s responses are received and processed by the scripts. The scripts extract the relevant information from the generated text, including the gene’s official name, function summary, evaluative comments for each criterion, and the corresponding scores. The extracted data is then stored in a structured format for further analysis and interpretation. This automated process allows for the efficient prioritization of many candidate genes without the need for manual prompt generation and response handling.

### Benchmarking experiments

#### Manual prompting

To assess the consistency and reproducibility of the gene prioritization results, we conducted benchmarking experiments comparing manual and automated prompting approaches. For manual prompting, we selected three different sites around the world: the United States, Thailand, and Qatar. At each site, a researcher manually submitted the same set of prompts to GPT-4 using the OpenAI web interface. The prompts were submitted in triplicate to evaluate the consistency of the model’s responses. The manual prompting process involved copying and pasting the predefined prompts into the GPT-4 interface, replacing the placeholder gene symbol with the actual gene of interest. The researchers then recorded the model’s responses, including the gene’s official name, function summary, evaluative comments, and scores for each criterion.

### API-based automated prompting

For the API-based automated prompting, we utilized the Python scripts developed for automating the candidate gene prioritization workflow. The scripts were configured to generate prompts for the same set of genes used in the manual prompting experiments. The automated prompts were submitted to GPT-4 via the OpenAI API, and the model’s responses were automatically processed and stored by the scripts. To assess the consistency and scalability of the automated approach, we performed the automated prompting experiments in triplicate and also increased the number of replicates to five. This allowed us to evaluate the reproducibility of the results and the potential for handling a larger number of candidate genes.

### Specific prompt used for benchmarking

For the benchmarking experiments, we used a specific prompt that assessed the gene’s relevance to various aspects of immune biology and its potential as a biomarker or therapeutic target, as detailed above and in supplementary methods. To ensure reproducibility and consistency across all experiments, we disabled web search capabilities in the ChatGPT interface and relied solely on the model’s pre-trained knowledge base, matching the API’s inherent functionality. This controlled knowledge environment ensured that all responses, whether through manual prompting or API calls, were based on the same knowledge cutoff date.

### Application of the automated candidate gene prioritization workflow for sepsis monitoring

#### Selection of candidate genes for prioritization

To demonstrate the scalability and potential clinical application of our automated candidate gene prioritization approach, we focused on developing a targeted assay for monitoring patients with sepsis. We selected the entire BloodGen3 repertoire, consisting of 382 modules, as the candidate gene pool for this use case.

#### Customization of prompts for sepsis-specific prioritization

We customized the prompt used in the automated prioritization workflow to focus on sepsis-specific criteria. The modified prompt assessed each gene’s relevance to sepsis pathogenesis, host immune response, organ dysfunction, biomarker potential, and therapeutic implications. The customized prompt was designed as follows:

For the Gene [input = official Gene Symbol]

1. Provide the gene’s official name
2. Provide a brief summary of the gene’s function.
3. Give each of the following statements a score from 0 to 10, with 0 indicating no evidence and 10 indicating very strong evidence:
4. The gene is associated with the pathogenesis of sepsis. Score: Based on evidence of the gene’s involvement in the biological processes underlying sepsis, including but not limited to its role in the dysregulated host response to infection, organ dysfunction, or sepsis-related complications.
5. The gene is associated with the host immune response in sepsis. Score: Based on evidence of the gene’s involvement in the immune response during sepsis, including but not limited to its role in innate or adaptive immunity, inflammation, or immunosuppression.
6. The gene is associated with sepsis-related organ dysfunction. Score: Based on evidence of the gene’s involvement in the development or progression of organ dysfunction in sepsis, including but not limited to its role in cardiovascular, respiratory, renal, hepatic, or neurological dysfunction.
7. The gene is relevant to circulating leukocytes immune biology in sepsis. Score: Based on evidence linking the gene to the development, function, or regulation of circulating leukocytes in the context of sepsis, including impacts on leukocyte differentiation, activation, signaling, or effector functions.
8. The gene or its products are currently being used as a biomarker for sepsis in clinical settings. Score: Based on evidence of the gene or its products’ application as biomarkers for diagnosis, prognosis, or monitoring of sepsis in clinical settings, with a focus on their validated use and acceptance in medical practice.
9. The gene has potential value as a blood transcriptional biomarker for sepsis. Score: Based on evidence supporting the gene’s expression patterns in blood cells as reflective of sepsis or its severity, considering both current research findings and potential for future clinical utility.
10. The gene is a known drug target for sepsis treatment. Score: Based on evidence of the gene or its encoded protein serving as a target for therapeutic intervention in sepsis, including approved drugs targeting this gene, compounds in clinical trials, or promising preclinical studies.
11. The gene is therapeutically relevant for managing sepsis or its complications. Score: Based on evidence linking the gene to the management or treatment of sepsis or its associated complications, including its role as a potential target for adjunctive therapies or personalized treatment strategies. Scoring criteria:

0 - No evidence found 1-3 - Very limited evidence 4-6 - Some evidence, but needs validation or is limited to certain conditions 7-8 - Good evidence 9-10 - Strong evidence

This prompt was integrated into the Python scripts that communicate with GPT-4 via the OpenAI and Microsoft Azure APIs, ensuring that the prioritization process was tailored to the specific requirements of sepsis monitoring.

### Execution of the automated prioritization workflow

We executed the automated candidate gene prioritization workflow using the customized prompts for sepsis monitoring. The Python scripts generated prompts for each candidate gene, incorporating the gene symbol and the predefined sepsis-specific criteria. The scripts then communicated with GPT-4 via the APIs, sending the prompts and receiving the model’s responses. The scripts extracted the relevant information from the generated text, including the gene’s official name, function summary, evaluative comments, and scores for each criterion.

### Utilization of LLMs for manuscript preparation

In addition to our primary research, we explored the potential of LLMs in manuscript preparation. Claude (Anthropic) was utilized under close human supervision to assist in drafting and editing, using our previous publication (18), study data, and key findings as context. All content was thoroughly reviewed and validated by the human authors. This application aligns with our broader exploration of GenAI in genomic research

### Code availability and deployment

Our research resources are readily available to the scientific community. An interactive web application has been developed using Streamlit (v1.30) for user-friendly access, which can be found at https://genellm.streamlit.app/. This application allows users to leverage our automated gene prioritization workflow through a user-friendly interface.

Key features of the web application include:

1. Personalized API Integration: Users can input their own API keys to initiate a session, ensuring secure and individualized access to the GPT-4 model.
2. Flexible Query Options: The application supports querying either a single gene or a list of genes, accommodating various research needs.
3. Parameter Tuning: Users can adjust various parameters to customize the prioritization process according to their specific requirements.
4. Interactive Supplementary Material: We have included interactive use cases that guide users through the application’s functionality, demonstrating its potential applications in different research scenarios.
5. Results Visualization: The application provides clear, easy-to-interpret visualizations of the prioritization results, enhancing data interpretation.

This web application not only makes our tool more accessible to researchers without extensive programming experience but also allows for rapid, iterative exploration of gene prioritization hypotheses. It represents a significant step towards making LLM-based gene prioritization a more widely available and user-friendly process in genomics research. The source code that underpins our tools is hosted on GitHub and is open for public use at https://github.com/taushifkhan/cd2k_llmWorkShop.git.

## RESULTS

### Development and Validation of a Python-based Framework for GenAI-Driven Gene Prioritization

Our study successfully developed and implemented a comprehensive framework for automated candidate gene prioritization, centered around custom Python scripts that interface with GenAI APIs. This technical achievement, utilizing GPT-4 via the OpenAI and Microsoft Azure APIs, represents a significant step in integrating advanced language models into bioinformatics workflows.

The Python scripts we developed not only automate the communication with LLMs but also handle complex tasks such as prompt generation, response processing, and data organization. This automation streamlines the gene prioritization process, making it more efficient and reproducible. We demonstrated the effectiveness of our framework through two key applications: (A) Benchmarking on interferon-associated modules: We initially focused on prioritizing genes associated with interferon responses in module M10.1 of the BloodGen3 repertoire. This served as a proof-of-concept and allowed us to fine-tune our approach. (B) Scaling to a larger gene set for sepsis analysis: To demonstrate the scalability of our framework, we applied it to a much larger gene set in the context of sepsis. This application showcases the framework’s ability to handle more complex, disease-specific prioritization tasks.

These applications highlight how our GenAI-based approach, powered by carefully crafted Python scripts, can be tailored to specific biological themes and scaled to address broader research questions, potentially accelerating the identification of key genes for targeted assay development across various contexts.

### Successful automation of the candidate gene prioritization workflow

The developed Python scripts successfully automated the candidate gene prioritization workflow using GPT-4 via the OpenAI and Microsoft Azure APIs. The scripts were able to generate the necessary prompts based on predefined templates and input parameters, including constructing prompts for gene scoring and prioritization, and incorporating specific gene symbols and other relevant information. The scripts efficiently handled the communication with the chosen API by sending the generated prompts to GPT-4 and retrieving the model’s responses. The returned data was processed and the relevant information, such as the gene’s official name, function summary, evaluative comments, and scores for each criterion, was extracted from the generated text.The extracted information was then stored in a structured format using Pandas DataFrames and stored in csv format for future use, facilitating further analysis and processing. The scripts organized the data in a way that streamlined the subsequent prioritization and interpretation of the results.

Overall, the developed Python scripts successfully abstracted away the complexities of API communication and data handling, allowing researchers to focus on other aspects of the prioritization and assay development process. The automation of the workflow using GPT-4 via the OpenAI and Microsoft Azure APIs demonstrated the feasibility and efficiency of integrating LLMs into the candidate gene prioritization pipeline.

### Effective prioritization of candidate genes in module M10.1 using GPT-4

Our testing focused on module M10.1, an interferon (IFN) response-associated module from the BloodGen3 repertoire (10). This module, one of six modules belonging to the A28 aggregate, all of which are functionally associated with IFN responses, comprises 21 genes (LAP3,OAS2,DDX58,PARP12,FBXO6,GBP4,GBP1,SAMD9L,SCO2,IFIT2,IFI35,DDX60,PAR P14,IFIH1,IRF7,TRIM22,ZBP1,STAT1,IFIT5,DHX58,GBP5).

The automated candidate gene prioritization process using GPT-4 and the designed prompts effectively generated informative and structured data for each gene in module M10.1. GPT-4 successfully provided the official gene name, a concise function summary, and evaluative comments for each specified criterion, allowing for a quantitative assessment of the evidence supporting a gene’s relevance to IFN responses, therapeutic potential, and other key aspects. **Figure 1** showcases a typical output generated by our automation script, which includes the top-scoring genes from module M10.1 (those with total scores above the 75th percentile) and their associated scores across eight evaluated criteria. The genes were selected based on their total score, calculated as the sum of the scores in each field. This structured output enables easy analysis and interpretation of the prioritization results.

**Figure 1.**
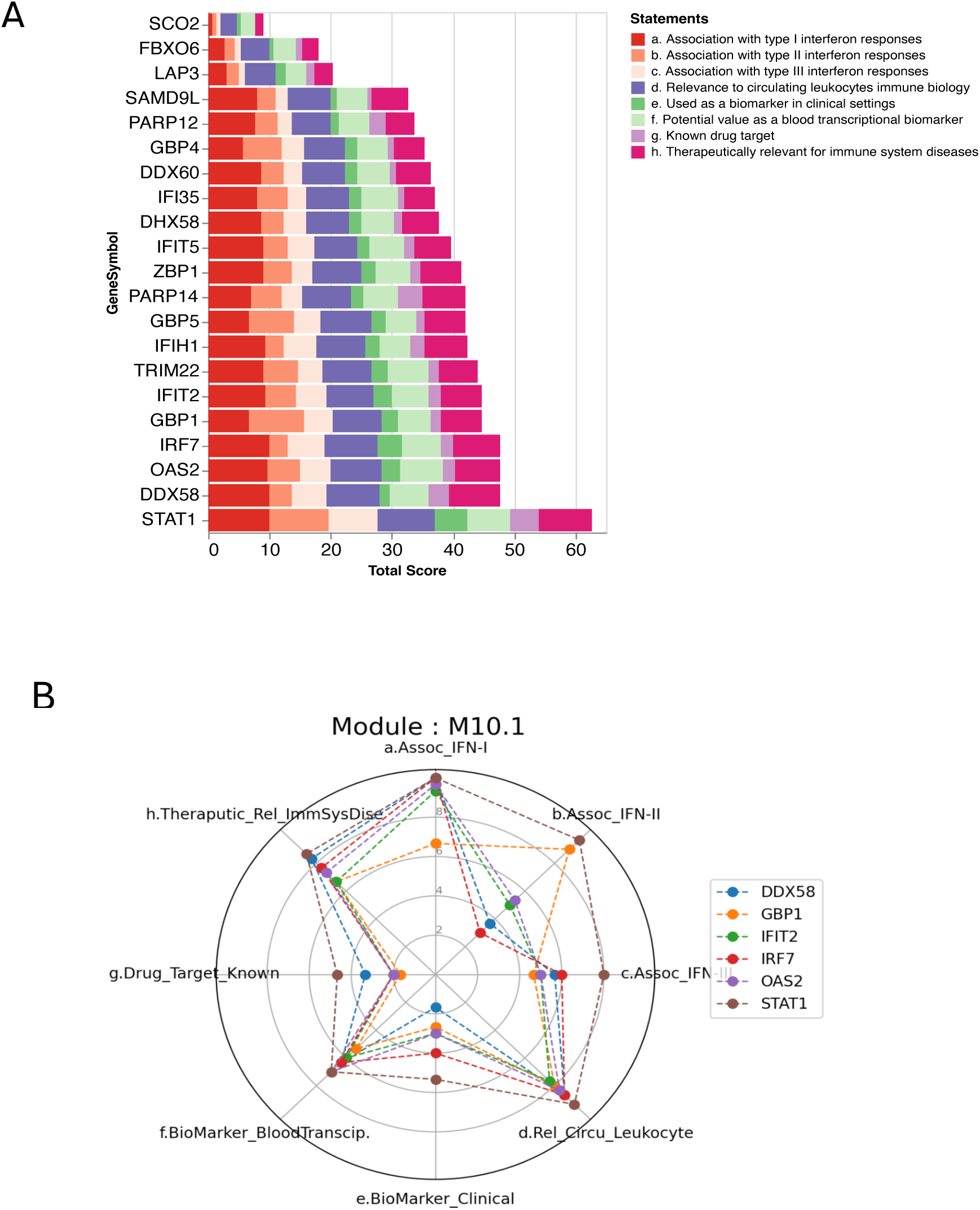
Automated candidate gene prioritization for module M10.1 using GPT-4. (A) Cumulative scores for all genes comprising the interferon response-associated module M10.1 from the BloodGen3 repertoire. The scores were generated using the GPT-4-based automated prioritization workflow, which evaluates each gene’s relevance to various criteria, such as association with interferon responses, potential as biomarkers, and therapeutic relevance. The cumulative score for each gene is calculated by summing its individual scores across all eight evaluated criteria (a-h), with a maximum possible score of 80 (8 criteria * 10 points each). (B) Spider graph displaying the scores of the top candidate genes across eight evaluated statements: (a) association with type I interferon, (b) association with type II interferon, (c) association with type III interferon, (d) relevance to circulating leukocytes, (e) current use as a clinical biomarker, (f) potential value as a clinical biomarker, (g) known as a drug target, and (h) therapeutic relevance for immune diseases. The scores range from 0 to 10, with higher scores indicating stronger evidence supporting the gene’s relevance to the corresponding statement.

Overall, the automated candidate gene prioritization process using GPT-4 effectively integrated LLMs into the workflow, generating informative and structured data for each gene in module M10.1. This approach enabled efficient prioritization based on the genes’ relevance to IFN responses and other key criteria, demonstrating the potential of LLMs in streamlining the gene prioritization process.

### Comparison of manual and automated prompting results

To validate the effectiveness and reliability of our API-based automated prompting approach, we conducted benchmarking experiments comparing its performance with the manual prompting method used in our previous study (18). These experiments aimed to assess the consistency and reproducibility of the gene prioritization results obtained using the two approaches and to evaluate the scalability of the automated method when dealing with a larger number of candidate genes.

The benchmarking experiments comparing manual and automated prompting approaches demonstrated reasonably high consistency and reproducibility in the gene prioritization results (**Figure 2**). Correlation analyses of the scores generated by GPT-4 revealed Pearson correlation coefficient values greater than 0.7 across all comparisons, indicating a strong positive relationship between the results obtained through different methods and sites (**Figure 2A**). Higher levels of correlation were observed within each approach, with coefficient values exceeding 0.8 for manual prompting experiments conducted at different sites (United States, Thailand, and Qatar) and values surpassing 0.9 for API-based automated prompting experiments performed with three and five replicates. These findings suggest that both manual and automated approaches yield consistent results when conducted in a controlled manner. Interestingly, the results generated manually in Qatar showed a higher level of correlation (coefficient > 0.85) with the results generated automatically (API) compared to the manual prompting results from the United States and Thailand (**Figure 2A**). This observation suggests that factors other than the prompting approach itself may influence the consistency of the results. Further investigation into the specific factors contributing to this difference is warranted.

**Figure 2.**
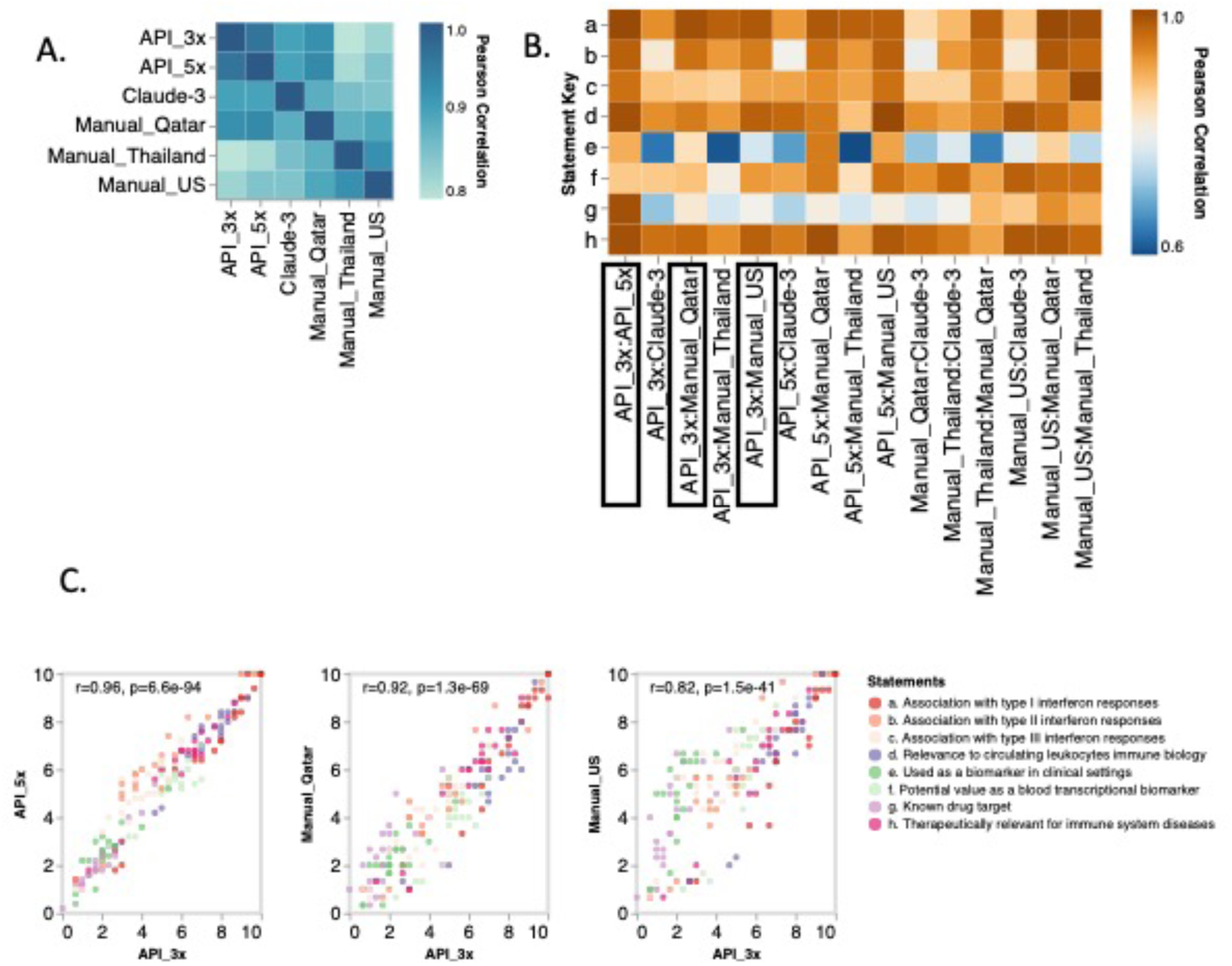
Consistency and reproducibility of gene prioritization results across manual and automated prompting approaches. (A) Heatmap displaying Pearson correlation coefficients between gene prioritization scores obtained using different methods (Claude-3, API 3x, API 5x, and manual prompting at three sites: Qatar, Thailand, and the United States). Higher correlation coefficients (darker shades) indicate stronger agreement between the results generated by the corresponding methods. (B) Heatmap showing Pearson correlation coefficients for individual statements assessed during gene prioritization. The heatmap reveals lower correlations (blue shades) for statements "e" (used as a clinical biomarker) and "g" (known as a drug target), suggesting potential variability in the evaluation of these specific criteria across different prompting approaches. (C) Scatter plots comparing gene prioritization scores obtained using the API-based automated approach with 3 replicates (API_3x) against scores from the API-based approach with 5 replicates (API_5x), manual prompting in the United States (Manual_US), and manual prompting in Qatar (Manual_Qatar). The high R-squared values indicate strong concordance between the scores generated by the automated and manual approaches. These plots are representative of the pairwise comparisons performed in the study.

The heatmap in **Figure 2B** reveals lower correlations for individual statements "e" (used as a clinical biomarker) and "g" (known as a drug target), suggesting potential variability in the evaluation of these specific criteria across different prompting approaches. This variability may be attributed to the complexity and specificity of the information required to assess these statements accurately. Scatter plots comparing gene prioritization scores obtained using the API-based automated approach with 3 replicates (API_3x) against scores from the API-based approach with 5 replicates (API_5x), manual prompting in the United States (Manual_US), and manual prompting in Qatar (Manual_Qatar) are presented in **Figure 2C**. The high R-squared values indicate strong concordance between the scores generated by the automated and manual approaches, further supporting the consistency and reproducibility of the prioritization results.

The API-based automation of the candidate gene prioritization workflow using GPT-4 offered several advantages over the manual prompting approach, including increased efficiency, reduced human error, and improved scalability. The automated scripts streamlined the process by efficiently generating prompts, extracting relevant information, and organizing the data into a structured format. The benchmarking experiments validated the effectiveness, reliability, and scalability of the API-based automated approach, highlighting its potential for application in larger-scale prioritization tasks. The strong correlations observed between the results obtained through manual and automated approaches, as well as across different sites, underscore the robustness and potential for widespread adoption of this methodology in the field of gene prioritization.

### Scalability of the automated candidate gene prioritization approach and its application in developing a targeted assay for monitoring sepsis

To demonstrate the scalability of our automated candidate gene prioritization approach, we performed a prioritization run across the entire BloodGen3 repertoire, which consists of 382 modules encompassing 11462 genes. After removing hypothetical and redundant genes, 10824 genes were analyzed for response. As a use case, we focused on developing a disease-specific targeted assay for monitoring patients with sepsis, a life-threatening organ dysfunction caused by a dysregulated host response to infection (20). Early detection and intervention are crucial for improving patient outcomes, and developing a targeted assay for sepsis could aid in the rapid identification of high-risk patients and guide personalized treatment strategies (21).

Our automated prioritization approach successfully identified sepsis-associated genes across the BloodGen3 repertoire, with distinct gene clusters emerging based on their response patterns to the evaluated criteria **(Figure 3A)**. Analysis of score distributions across all eight criteria revealed biologically meaningful right-skewed patterns, particularly pronounced in criteria related to clinical biomarkers and drug targets (Supplementary Figure S1). This skewness reflects real-world expectations, where only a subset of genes would have strong documented evidence for specific functions or clinical applications. After filtering for genes with scores > 5 in at least one criterion, the remaining distribution patterns helped identify candidates with substantial evidence across multiple criteria, supporting our gene prioritization strategy. The analysis revealed the proportion and distribution of sepsis-associated genes within module aggregates **(Figure 3B)** and individual modules **(Figure 3C)**, providing valuable insights for the development of targeted assays. The analysis of the distribution of sepsis-associated genes across the BloodGen3 module aggregates (**Figure 3B**) shows a striking concordance with the functional annotations attributed to these aggregates by Altman et al. (10), as depicted in the image provided.

**Figure 3.**
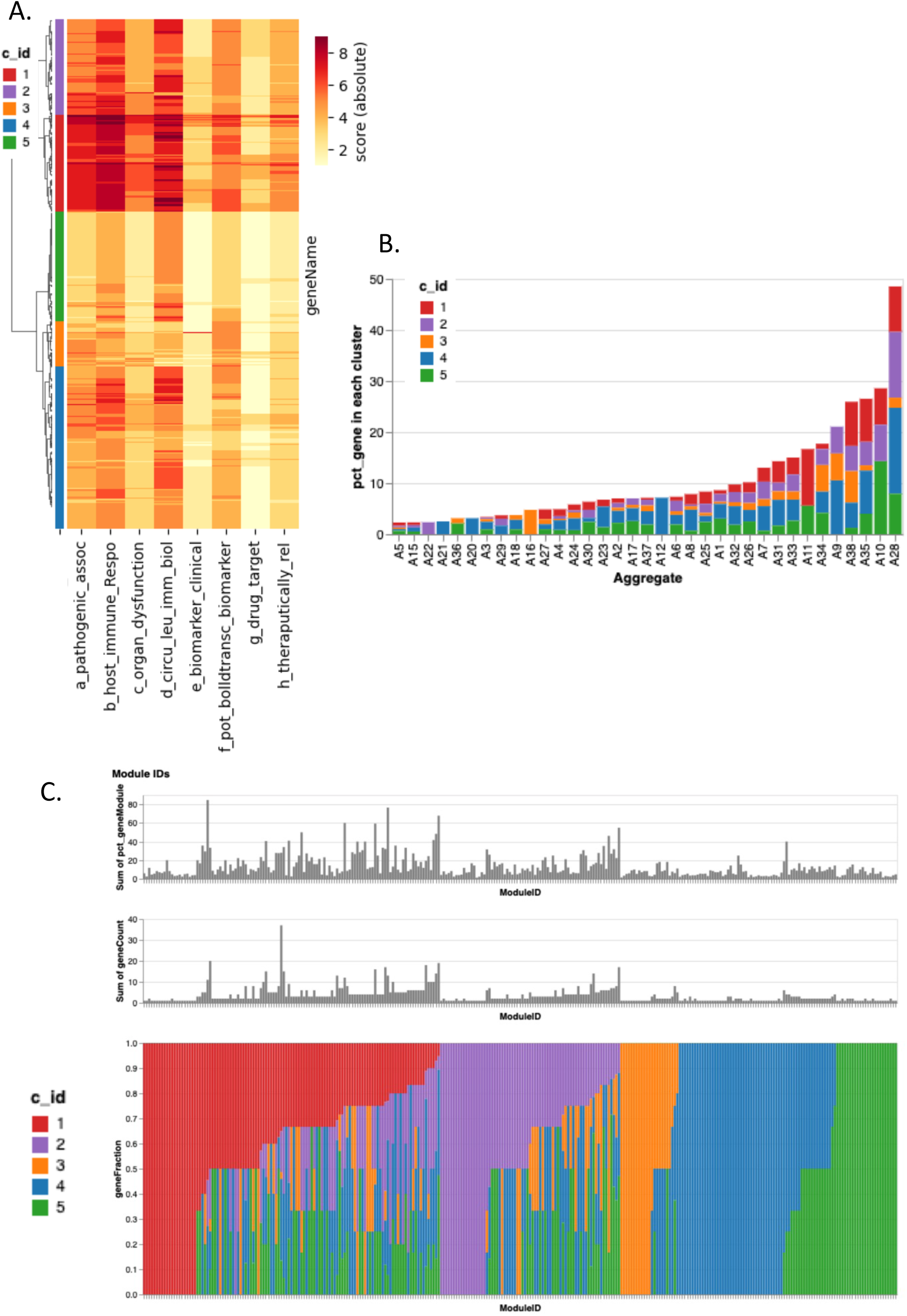
Application of the automated candidate gene prioritization workflow for sepsis monitoring. Figure 3. [A] Hierarchical Clustering of Sepsis-Related Genes Based on Response Scores : This figure presents a hierarchical clustering analysis of 1,070 genes (rows) filtered based on their relevance to sepsis-related queries (i.e genes with score > 5 in at least one of the sepsis related query). The heatmap displays the absolute response scores for each gene, with colors ranging from white (low scores) to orange to red (high scores). Genes are grouped into clusters based on their overall response patterns, as indicated by the color-coded track along the hierarchical clustering tree. [B] Percentage of Gene in each cluster across Module Aggregates: Bar plot shows distribution of sepsis related genes (n=1070) across pre-defined BloodGen3 Module Aggregates (= 35 [out of 38]). Each bar shows percentage of genes found in each Module Aggregate, as well proportionality of gene clusters with same color code as shown in A. [C] Distribution of Gene Membership Across Clusters in Different Modules: Stacked bar plot visualizes the distribution of gene membership across different modules related to sepsis, with each module represented along the x-axis (total modules = 297 [out of 382 Modules in BloodGen3]). The y-axis displays the fraction of member genes within each cluster, represented by stacked color bars corresponding to each cluster. The calculation for the representation of cluster members in each module is performed by dividing the number of genes in a cluster for a specific ModuleID by the total number of genes in that respective ModuleID. The top panel of the figure provides additional context by showing the total number and percentage of genes in each respective module, helping to understand the scale and relevance of each module’s gene population.

The module aggregates with the highest proportions of sepsis-associated genes identified by our automated workflow include A35 (inflammation, neutrophil activation), A28 (interferon responses), A32-A34 (cytokines/chemokines, inflammation, leukocyte activation), and A8 (monocytes). This aligns well with the known pathophysiology of sepsis, which is characterized by a dysregulated host immune response involving excessive inflammation, neutrophil and monocyte activation, and cytokine production (20). Interferon signaling has also been implicated in the pathogenesis of sepsis (22).

Conversely, module aggregates associated with adaptive immunity, such as A1-A3 (T cells, B cells) and A17 (antigen presentation), show a lower representation of sepsis-associated genes. This is consistent with the notion that sepsis is primarily driven by innate immune responses, while adaptive immunity may be suppressed in the acute phase of the disease (23).

The concordance between the distribution of sepsis-associated genes identified by our automated prioritization approach and the known functional associations of the BloodGen3 module aggregates provides a strong biological validation of our findings. It demonstrates the ability of our workflow to capture relevant disease-associated gene signatures and highlights its potential utility in designing targeted assays for sepsis monitoring and management.

Notably, the entire prioritization run, which involved processing all genes in the BloodGen3 repertoire across 8 statements took approximately 3 days to complete and incurred a cost of about $200. This demonstrates the feasibility and cost-effectiveness of applying our automated approach to large-scale gene prioritization tasks.The results obtained from this prioritization run highlight the potential of our automated workflow to efficiently identify disease-associated genes across extensive gene repertoires. The insights gained from this analysis can guide the development of targeted assays for sepsis monitoring, focusing on the most relevant gene clusters and modules.

## DISCUSSION

This study presents the development and benchmarking of an automated workflow for prioritizing candidate genes, utilizing large language models (LLMs) like GPT-4, to enhance the identification of promising genes for targeted assay development. The method involves Python scripts that interact with GPT-4 through OpenAI and Microsoft Azure APIs, automating the creation and submission of prompts and the extraction of relevant information. To ensure reproducibility and consistency in model responses, we implemented standardized API parameter settings, including temperature and sampling configurations. Customizable prompts were designed to evaluate candidate genes based on criteria such as association with biological processes, biomarker potential, and therapeutic implications, tailored for specific diseases or processes. By incorporating sepsis-specific criteria into the prompts and considering technical requirements for targeted assay development, we demonstrated the flexibility and scalability of our automated candidate gene prioritization approach. This showcased the potential for adapting the workflow to design targeted assays for various diseases, such as sepsis, based on the specific biological and clinical context.

To validate the effectiveness and reliability of the API-based automated prompting approach, we conducted benchmarking experiments comparing its performance with the manual prompting method. While API usage is generally considered best practice for large-scale data tasks, this comparison provided valuable insights into the consistency and reproducibility advantages of our automated approach. These experiments aimed to assess the consistency and reproducibility of the gene prioritization results obtained using the two approaches, as well as to evaluate the scalability of the automated method when dealing with a larger number of candidate genes. By demonstrating the comparability of the API-based approach to the previously established manual workflow, we sought to provide a strong foundation for the adoption of automated LLM-based gene prioritization in the design of targeted assays for various biological and clinical applications. The workflow was applied to prioritize genes from the BloodGen3 repertoire, comprising 382 modules, for sepsis assay development. This study underscores the potential of incorporating LLMs into gene prioritization, offering a more efficient and systematic way to identify candidates for targeted assays, adaptable to various biological and disease contexts.

The application of large language models (LLMs) in biomedical research has recently gained attention, with several studies exploring their potential in various tasks, such as disease gene prioritization. Kim et al. (24) conducted a comprehensive evaluation of five LLMs, including GPT and Llama2 series, for phenotype-driven gene prioritization in rare genetic disorder diagnosis. While their findings revealed that the best-performing LLM, GPT-4, achieved an accuracy of 16.0%, it still lagging behind traditional bioinformatics tools. However, they observed that prediction accuracy increased with the parameter/model size, highlighting the potential for further improvements as LLMs continue to evolve.

In the context of knowledge-driven gene prioritization, several approaches have been proposed that leverage various data sources and algorithms. For example, Emad et al. (25) developed a method called ProGENI that utilizes prior knowledge of protein-protein and genetic interactions, along with gene expression data, to prioritize genes associated with drug response. Their results demonstrated that knowledge-guided prioritization outperformed other methods and provided new insights into the mechanisms of drug resistance.

Another notable approach is the Monarch Initiative (26), which integrates open data at a global scale to bridge the gap between genetic variations, environmental determinants, and phenotypic outcomes. The Monarch App combines data about genes, phenotypes, and diseases across species, enabling advanced analysis tools for variant prioritization, deep phenotyping, and patient profile-matching. Interestingly, the Monarch Initiative has also explored the integration of LLMs, such as OpenAI’s ChatGPT, to increase the reliability of its responses about phenotypic data. While these studies showcase the potential of knowledge-driven gene prioritization, they often rely on curated databases and predefined feature sets. In contrast, our approach leverages the vast knowledge base and natural language understanding capabilities of LLMs to assess the relevance of candidate genes based on a wide range of criteria and the most up-to-date information available in the biomedical literature.

It is important to note that our work focuses on the prioritization aspect of the gene discovery process, and that the final selection of candidate genes still requires additional steps and manual intervention. Due to the large scale of our study, performing manual fact-checking on the scoring justifications, as we have done in previous work, would be highly laborious or even infeasible, which is another notable limitation. Additionally, we observed some inconsistencies in the scoring, as illustrated in Figure 2B and discussed in the ’Comparison of manual and automated prompting results’ section, which, coupled with the lack of extensive manual validation, suggests that our work serves more as a proof of concept at this stage, exploring the current capabilities of LLMs in this context. It is also worth noting that the final gene selection process should consider other factors, such as gene expression levels and consistency across reference disease cohorts.

Another limitation of our work is that it was demonstrated on a specific use case, focusing on the prioritization of candidate genes for sepsis monitoring. The prompts could however be readily adapted to other disease settings, either manually or using LLMs for prompt engineering. While our current framework demonstrates the utility of basic prompting strategies for gene prioritization, we acknowledge the potential for incorporating more advanced Natural Language Processing (NLP) techniques. Recent developments in prompt engineering, such as in-context learning, chain of thought prompting, tree of thought, and Retrieval-Augmented Generation (RAG), offer promising avenues for enhancement. Our framework’s modular design allows for the integration of such techniques, though careful evaluation will be needed to balance increased complexity against practical utility in gene prioritization tasks. Future iterations of this work could particularly benefit from RAG strategies to incorporate up-to-date scientific literature, potentially improving the accuracy and comprehensiveness of gene assessments. However, the current implementation prioritizes accessibility and ease of use while maintaining effective performance for gene prioritization tasks.

In conclusion, our study demonstrates the potential of integrating large language models (LLMs) into the candidate gene prioritization process, offering a more efficient and systematic approach to identifying promising candidates for targeted assay development. By leveraging the vast knowledge base and natural language understanding capabilities of LLMs, our automated workflow can assess the relevance of candidate genes based on a wide range of criteria and the most up-to-date information available in the biomedical literature. The flexibility and adaptability of our approach opens new avenues for the design of targeted assays for diverse applications, such as disease monitoring, treatment response assessment, and the identification of novel therapeutic targets. As LLMs continue to evolve and improve, we anticipate that their integration into biomedical research workflows will become increasingly valuable, enabling researchers to navigate the ever-growing body of knowledge more effectively and efficiently. Future work should focus on further validating and refining our approach across different biological contexts and disease states, as well as exploring the integration of additional data sources and analysis techniques to enhance the accuracy and robustness of the prioritization process. Moreover, the development of standardized protocols and best practices for the use of LLMs in biomedical research will be crucial to ensure the reproducibility and comparability of results across studies. The integration of LLMs into the candidate gene prioritization process has significant implications for the development of targeted assays, particularly in the context of large-scale immune monitoring and research in low-resource settings. By streamlining the identification of the most promising candidate genes, our approach can facilitate the design of cost-effective and targeted assays that focus on the most relevant biomarkers for a given disease or biological process.

